# The positive role of noise for information acquisition in biological signaling pathways

**DOI:** 10.1101/762989

**Authors:** Eugenio Azpeitia, Andreas Wagner

## Abstract

All living systems acquire information about their environment. At the cellular level, they do so through signaling pathways, which rely on interactions between molecules that detect and transmit the presence of an extracellular cue or signal to the cell’s interior. Such interactions are inherently stochastic and thus noisy. In classical information theory, a noisy communication channel degrades the amount of transmissible information relative to a noise-free channel. For this reason, one would expect that the kinetic parameters that determine a pathway’s operation minimize noise. We show that this is not the case under a wide range of biologically sensible parameter values. Specifically, we perform computational simulations of simple signaling systems, which show that a noisy molecular interaction dynamics is a necessary condition for information acquisition. Moreover, we show that optimal information acquisition, where a system reacts most sensitively to changes in the environment, can be obtained close to the maximal attainable level of noise in the system. Our work highlights the positive role that noise can have in biological information processing.

**Author summary:** The acquisition of information is fundamental for living systems, because the decisions they take based on such information directly affect survival and reproduction. The molecular mechanisms used by cells to acquire information are signaling pathways. The molecular interactions of signaling pathways, such as the binding of a signal to a receptor, are by nature noisy. This is important, because noise disrupts information. Hence, to maximize the acquisition of information, signaling pathways should minimize the noise of their molecular interactions. Here we show that a noisy dynamic of the molecular interactions can improve the acquisition of information, and that the maximal capacity to acquire information can be obtained with a close-to-maximal level of noise in a signaling pathway. Thus, contrary to expectations, noise can improve the acquisition of information in living systems.

## Introduction

Information about the environment is fundamental when living organisms make decisions that affect their survival and reproduction (1). For example, microbes adjust their growth in response to nutrient concentrations, animals flee in response to predators, and plants synthesize defense chemicals in response to herbivores.

At the cellular level, signaling pathways are the main molecular mechanism by which organisms acquire information. They typically detect the presence of a molecular signal or cue (2) about the environment through the binding of this molecule to a receptor. Once the signal has been detected, a chain of intermediary events transmits this information to the cell’s interior, where it ultimately regulates gene expression. Signaling pathways vary widely, including in their number of molecular interactions, signal and receptor affinities, the presence of feedback and feed-forward interactions, and the number of regulated genes (3,4). However, they all share some elementary processes, such as the reversible binding of molecules, which is necessary to detect a signal by a receptor, transmit its presence via effector molecules, for example through allosteric control of these molecules, and regulate gene expression through the binding of transcription factors to DNA.

Noise is present at all spatial and temporal scales of biological organization, from population dynamics to molecular interactions, including signaling pathways (5,6). The sources of noise in signaling pathways include random fluctuations in the concentration, movement, activity, and interactions of molecules (7–12). Classical information theory predicts that noise degrades the capability of a communication channel to transmit information. For example, in simple systems such as a binary or a symmetric information transmission channel, the maximum capacity of the channel to transmit information can only be realized in the absence of noise (13). In signaling pathways, noise transforms a stimulus, such as the concentration of a nutrient, into a distribution of outputs or responses. Any overlap between the response distributions produced by two different stimuli, such as two different signal concentrations, creates uncertainty about which stimuli produced which output (Fig 1a; 14). For this reason, noise also decreases the information acquisition capacity of signaling pathways.

**Fig 1.**
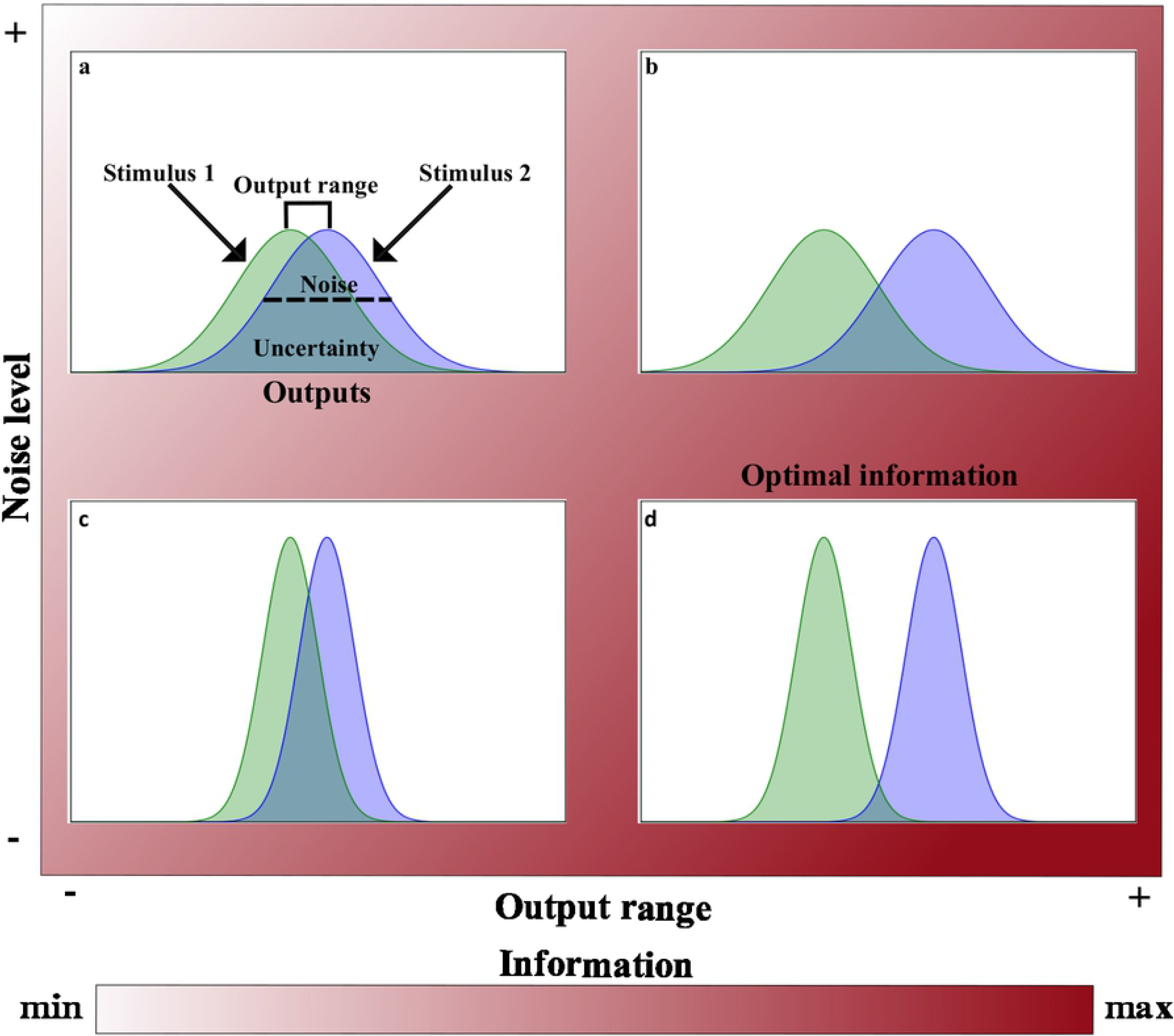
Relationship of acquired information with both noise and output range. Example of two hypothetical stimuli that produce different but overlapping response distributions (green and blue distributions). The amount of information acquired at different levels of noise (*y* axis) and with different output ranges (*x* axis) is indicated by the color bar. The black dashed line in (a) is a schematic representation of noise (i.e., the standard deviation of the response distributions, green and blue). The square bracket above the response distributions in (a) indicates the output range (i.e., the maximal difference of the mean values of the response distributions). Increasing the output range (b) and reducing the noise level (c) decrease the overlap (uncertainty) between response distributions observed in (a). (d) Acquired information is largest (maximal) when noise is minimized and the output range is maximized.

The overlap between response distributions can be reduced in two ways (11). First, the response distributions can be made more distinct by separating their means while preserving their dispersion (e.g., their standard deviation). This implies that the range of outputs produced by the stimuli will increase (Fig 1b). The second way to decrease the overlap between the response distributions is to decrease their dispersion, while keeping the mean constant. This is equivalent to reducing noise, which allows detecting the signal with increasing precision (Fig 1c). Thus, information acquisition is maximized when the output range is maximized and noise is minimized (Fig 1d).

Various mechanisms can either reduce noise or increase the output range to improve information acquisition (10,11,15–17). These include feedback loops and protein-protein interactions that reduce the level of noise (12,18,19), and increasing the number of molecules, which increase the output range (20,21). As a result, we know that noise – and thus also information acquisition – can be tuned within some limits (20,22–26). However, it is less clear how signaling pathways adjust their kinetic parameters, such as the association and dissociation rate of binding molecules, to minimize noise or increase the output range to respond efficiently to an environmental signal.

While a few studies have explored the effect of kinetic parameters on information transmission in signaling pathways, small gene networks, and gene expression systems (10,11,16,26), most of these studies did not explicitly model the molecular interactions involved in signaling. Therefore, they provide little intuition about why and how the kinetic properties of molecular processes affect information acquisition. To overcome this limitation, we use models that explicitly include all relevant molecular interactions and that do not make any a priori assumptions about statistical properties of noise. With these models, we analyze how the kinetic properties of the reversible binding interactions used by signaling pathways affect the relationship between noise, output range and information acquisition. First, we study the relationship between noise, output range and information in the reversible binding of two molecules that represent a signal and a receptor. We then analyze how information is transmitted in a chain of consecutive binding interactions. We then focus on information acquisition in gene regulation by the reversible binding of a TF to the DNA. Finally, we assemble all these components to help us understand information acquisition in a simple model of a linear signaling pathway. Our results show that, contrary to what is expected, under a broad range of biochemically sensible parameters, a noisy dynamic of the molecular interactions increases information acquisition in signaling pathways.

## Results

### The models

We study multiple models that represent either different fundamental steps of a signaling pathway or a complete pathway. All these models include an input or signal molecule *S* and an output *O* that conveys information about the signal’s value. We quantified (1) noise as the average standard deviation of the response distributions, (2) the output range as the maximal difference of the means of the response distributions, and (3) information as the mutual information between the signal and the output (see Methods). We estimate these quantities through at least 1000 stochastic simulations for each of *n* evenly distributed values of the number of signal molecules (*N_S_*) within the interval [*N_smax_*/*nN_smax_*].

In all our models, the signal is detected by reversibly binding of a molecule to either a receptor *R* or to a DNA binding site (*DNA_bs_*). Hence, all models contain at least one reversible binding interaction between molecules. We describe the affinity of two reversibly binding molecules with the equilibrium constant *K_eq_*(*M*)=*k_d_*/*k_a_*, where *k_d_* and *k_a_* represent the dissociation and association rate, respectively. The equilibrium constant represents the concentration of free signal molecules at which half of the receptors are bound to a signal molecule. As the equilibrium constant decreases, the concentration of signal molecules required to occupy 50% of the receptors decreases too. Hence, smaller *K_eq_* means higher affinity.

Throughout this paper, we will refer to low, intermediate and high affinities in the following sense. A low affinity refers to an equilibrium constant that is much higher than the maximal concentration of the signal. A high affinity refers to an equilibrium constant that is much lower than the maximal concentration of the signal. Finally, an intermediate affinity refers to an equilibrium constant that is between the minimal and the maximal concentration of the signal. In all our models we considered biologically meaningful values of all biochemical parameters (See Methods and S1-4 Tables).

### Reversible binding of molecules

We first study the reversible binding between two types of molecules, *S* and *R* that form *RS* complexes (Fig 2a). In this highly simplified model of an information transmission system, we consider the number of *RS* complexes as the output or response that conveys information about the presence of the signal *S*. Although this notation is suggestive of interactions between a signal (S) and a receptor (R), our framework below applies to any other reversible binding of two molecules that form a complex. However, for simplicity, we will refer to *R* molecules as receptors, and to *S* molecules as signal molecules.

**Fig 2.**
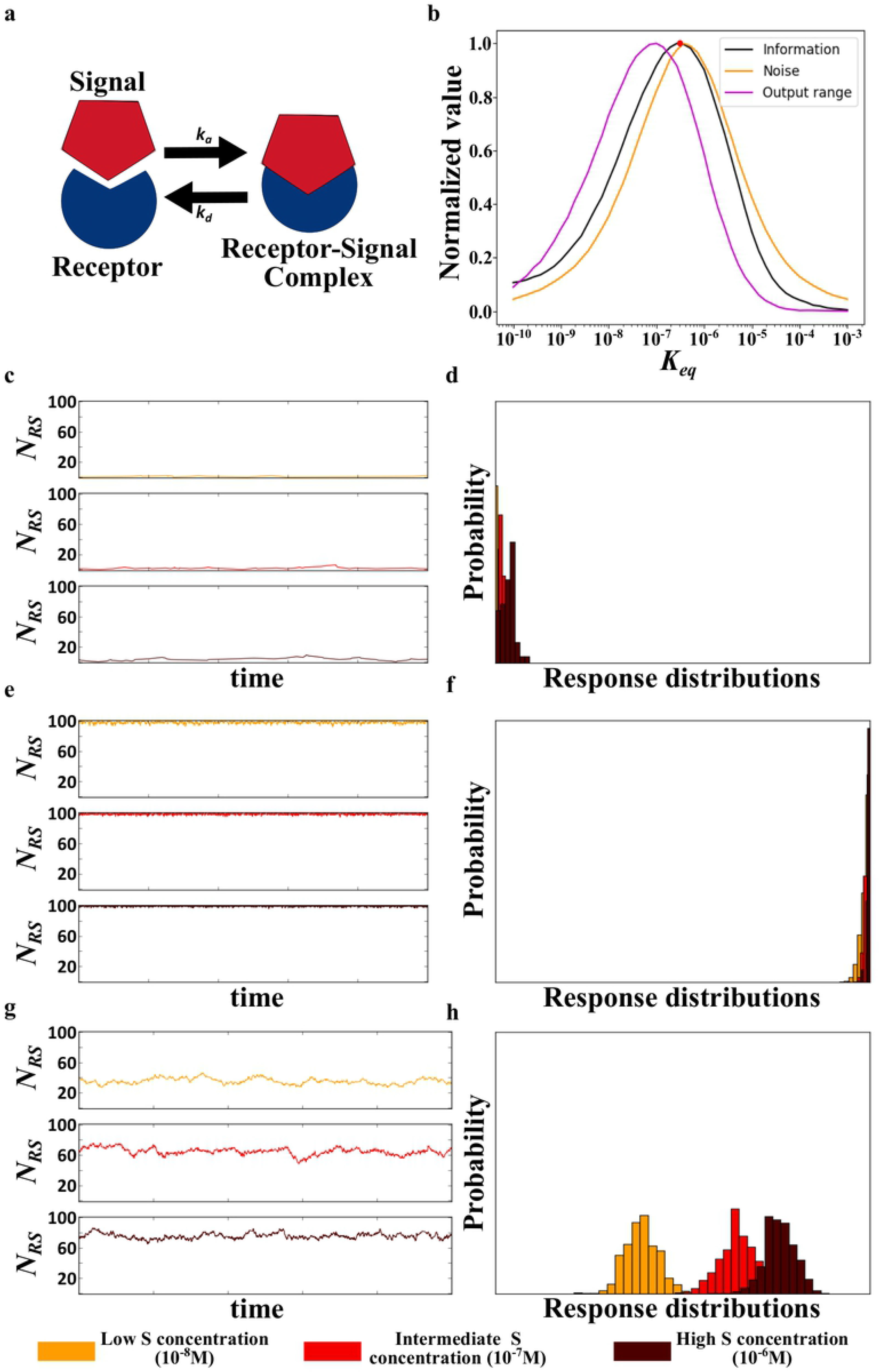
Noise, output range and information in the reversible binding of molecules. (a) Schematic representation of reversible binding involving a receptor and a signal as examples. *k_a_* and *k_d_* correspond to the association and dissociation rate, respectively. (b) Acquired information, output range, and noise for receptor-ligand binding at different affinity values (*K_eq_*). The red circle denotes the affinity at which mutual information between signal and output is optimized. Information, noise, and output range are normalized by their respective maximal values. Further panels show the system’s behavior at (c,d) low affinity (*K_eq_*=10^-5^), (e,f) high affinity (*K_eq_*=10^-9^), and (g,h) intermediate affinity (*K_eq_*=10^-7^;g-h). (c, e, g) show the temporal dynamic of the receptor-signal complexes (N_RS_) at three different concentrations of the signal *S*. (d, f, h) show response distributions at these signal concentrations (see color legend at the bottom of the figure).

We asked how noise, output range and information change with the affinity between receptor and signal molecules. For this analysis, we assumed that the concentration of the receptors is 10^-8^M and that the concentration of signal molecules lies within the range [10^-8^M,10^-6^M]. This means that the maximal number of signal molecules is greater than the total number of receptors. Our simulations allow us to distinguish three regimes as a function of affinity. First, when affinity is low, noise, output range and information are close to zero (Fig 2b). The reason is that few receptor-signal complexes form at low affinity, regardless of the signal concentration (S1 Fig). Consequently, the number of receptor-signal complexes (*N_RS_*) is close to zero for all values of the signal concentration (i.e., *N_RS_*➔0 for all *N_s_* as *K_eq_*➔∞; Fig 2c). For this reason, response distributions overlap greatly (Fig 2d), causing the output range and information to approach zero (Fig 2b). In addition, noise is also close to zero because the number of receptor-signal complexes fluctuates little (Fig 2c).

Second, when affinity is high, receptors are saturated at most or all signal concentrations (S1 Fig), such that the number of receptor-signal complexes is equal to the total number of receptors (i.e., *N_RS_*➔*N_R_* for all *N_S_* as *K_eq_*➔0; Fig 2e), and the output range approaches zero (Fig 2b). Noise approaches zero as well (Fig 2b), because the number of receptor-signal complexes barely fluctuates from its large value (Fig 2e), and because the overlap between response distributions is large (Fig 2f), acquired information approaches zero as well (Fig. 2b).

All this changes at intermediate affinities, where receptors can acquire information about the number of signal molecules, because receptors are no longer mainly saturated or unoccupied. Instead, the number of receptor-signal complexes fluctuates (Fig 2g). These fluctuations increase noise, but at the same time they permit that the mean number of receptor-signal complexes differs for different number of signal molecules. As a result, the output range increases (Fig 2b), which decreases the overlap between output distributions (Fig 2h), increasing the acquired information (Fig 2b). These observations show that a noisy signal-receptor binding dynamics can be beneficial when a receptor is to acquire information about a signal. Remarkably, the amount of acquired information is maximal when noise is close to its maximally possible value (Fig 2b). In sum, if information is acquired through reversible binding interactions, the binding kinetics that yield close to maximal noise also yield close to maximal information.

Next, we wondered if one can preserve the low noise of high and low affinity binding, while increasing the output range to maximize information acquisition. In doing so, we studied how changes in signal and receptor concentrations affect information, noise and output range. These concentrations, together with the affinity, completely determine the system’s behavior. We varied these concentrations in two different ways. First, we varied the concentrations of the receptors and signal molecules by identical amounts, which keep the ratio of receptors to signal molecules constant. Second, we only varied the concentration of the signal, which changes this ratio. In both cases, we found the same qualitative relationship between noise, output range, and information as before, as long as the maximal number of signal molecules is in excess of the number of receptors. In other words, efficient information acquisition requires high levels of noise and a high output range (Fig 3a-c; S2 Fig). The higher the signal and receptor concentrations are, the lower are the affinity values required for efficient information acquisition (Fig 3a). The reason is that a receptor’s affinity to its signal needs to decrease as signal concentration increases; otherwise receptors become saturated and no longer detect signal changes effectively.

**Fig 3.**
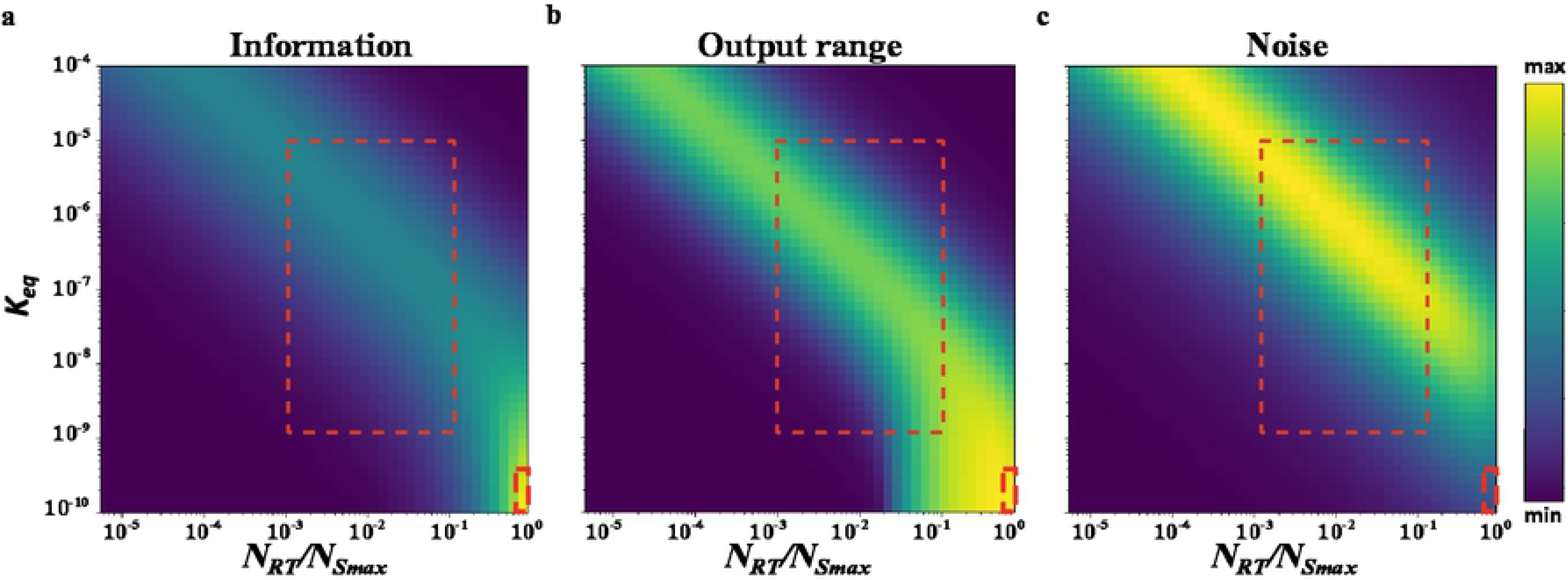
Acquired information, output range and noise at different signal concentrations. Contour plots of (a) information, (b) output range, and (c) noise at different receptor signal ratios *N_RT_/N_Smax_* (*x* axis), and at different affinities (*y* axis). (a-c) The large red-dashed rectangles circumscribe biologically common ranges of receptor-signal affinities ([10^-6^M,10^-9^M]) and *N_RT_/N_Smax_* ratios (*N_RT_*=10^-8^ and 10^-7^M≤*N_Smax_*≤10^-5^M). The small red-dashed rectangle circumscribes the region where *N_Smax_*=*N_RT_* and where the system is noise-free, reaches the maximally possible output range, and where information acquisition is ‘perfect’. Acquired information, output range and noise are plotted from minimally to maximally observed values, color-coded as indicated by the color bar.

The only scenario where low noise allows maximal information acquisition requires fewer signal molecules than receptors (Fig 3a-c, lower right corners small red rectangle). As an extreme case, one can think of a system with an infinitely large number of receptors, a finite number of signal molecules, and extremely high receptor-signal affinity. In such a system all signal molecules are bound to receptors. Because there are fewer signal molecules than receptors, the system effectively ‘counts’ the number of signal molecules through the number of receptor-signal complexes. Notice that experimentally measured affinity values between receptors and signals, are not extremely high. Instead, they are of the same order of magnitude as signal and receptor concentrations (11,28). In our simulations, these are the affinity values where high information acquisition entails high noise (Fig 3a-c big red rectangle), suggesting that biological system operate in the noisy regime. In sum, under biologically feasible conditions, high noise is necessary for (maximal) information acquisition.

### Consecutive reversible binding interactions

In a signaling pathway, the binding of a signal to a receptor is usually the first of a chain of reversible events. These events include the reversible modification of one or more intermediary signaling molecules, and they usually terminate in the reversible binding of transcriptional regulators, such as a transcription factors (TF), to DNA. TF-DNA binding differs from other signaling binding interactions because regulated genes have one or few copies in any one genome, and any one regulated gene harbors few – usually fewer than ten – TF-binding sites (29,30). In the simplest signaling pathways, signal-bound receptors can directly regulate transcription without intervening signaling steps (31).

To study how TF-DNA binding might affect information acquisition in such a pathway, we model two consecutive reversible binding interactions. They represent the formation of a receptor-signal complex, and the binding of this complex to a DNA binding site (*DNA_bs_*). We assumed that the concentration of receptor molecules is 10^-8^M, that the concentration of signal molecules lies in the interval [10^-8^M, 10^-6^M] and that a single DNA binding site mediates transcriptional regulation. The receptor-signal-DNA_bs_ complex represents the ultimate output of the system that harbors information about the signal.

We analyzed how the affinities of both the receptor to the signal (*K_eqR,S_*) and of the receptor-signal complex to DNA (*K_eqRS,D_*) affect the acquisition of information, output range and noise. As in the simpler two-molecule system, the receptor is able to detect different signal concentration at intermediary affinity values between the receptor and the signal, where the largest output ranges, which are necessary for the receptor to sense the signal, are produced with high levels of noise (Fig 4a-c).

**Fig 4.**
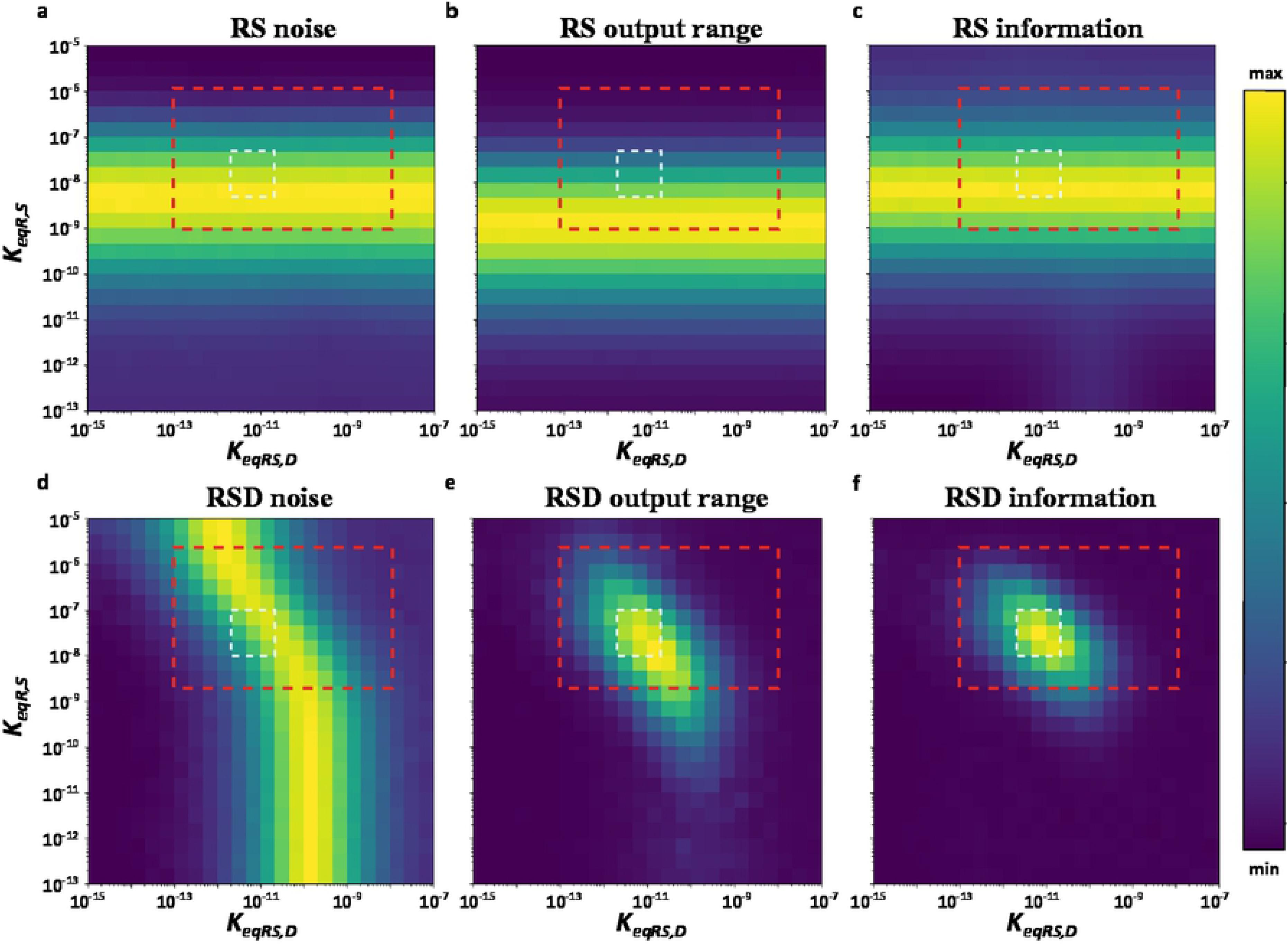
Information, and noise in a pair of reversible binding interactions. Contour plots of (a and d) noise, output range (b and e) and information acquisition (c and f) in the receptor-signal complex (*RS*; a-c) and in the receptor-signal-*DNA_bs_* complex (*RSD*; d-f) as a function of the affinities between both the receptor and the signal (*K_eq^R,S^_*), and the receptor-signal complex with the downstream molecule (*K_eqRS,D_*). Red-dashed rectangles circumscribe biologically sensible receptor-signal DNA affinities ([10^-8^M,10^-13^M]) and receptor signal affinities ([10^-6^M,10^-9^M]). White-dashed rectangles delineate the region of maximal information acquisition at the receptor-signal-*DNA_bs_* level. Acquired information, output range and noise are plotted from minimally to maximally observed values, color-coded as indicated by the color bar.

To subsequently transmit the information acquired by the receptor-signal complex to the receptor-signal-*DNA_bs_* complex, DNA binding needs to be subject to the same kind of fluctuations. Such fluctuations only occur at intermediary affinity values between the receptor-signal complex and the DNA, otherwise the DNA binding site is either almost always bound (saturated) or unbound by the receptor-signal molecules. The fluctuations in receptor-signal-DNA binding increase noise (Fig 4d), but they also lead to different probabilities of DNA binding for different concentrations of the receptor-signal complex, which increases the output range (Fig 4e). As a result, the acquisition of information increases (Fig 4f). In sum, information about a signal is obtained at intermediate values of both affinities (compare the white rectangles, indicating the region with maximal information at the receptor-signal-*DNA_bs_* level in Fig 4). We also note that the affinities leading to high information acquisition and high noise in our model are similar to experimentally measured affinities between receptors and signals, as well as between transcriptional regulators to DNA (Fig 4, large red rectangles). Repeating our analysis with up to ten DNA binding sites leads to the same conclusions (S3 Fig): A noisy dynamic is essential to acquire information.

### Gene expression system

At the end of signaling pathways stands the regulation of gene expression, which usually requires reversible binding (of a transcription factor to DNA), and additionally involves the synthesis and degradation of mRNA and protein. To find out whether the observed relationship between information, output range, and noise is similar in the presence of synthesis and degradation, we modeled the regulation of gene expression mediated by a transcription factor that reversibly binds to DNA. We assumed that a gene with a single DNA binding site drives transcription initiation, which occurs only when the binding site is bound by a transcription factor. In this case, mRNA molecules are transcribed at rate *k_x_*, and proteins are translated from the mRNA molecules at a rate *k*_2_. Both mRNA and protein molecules become degraded at rates, *d_x_* and *d*_2_, respectively (Fig 5a). We considered the number of TF molecules as the signal, and the number of protein molecules *N_P_* as the output or response.

**Fig 5.**
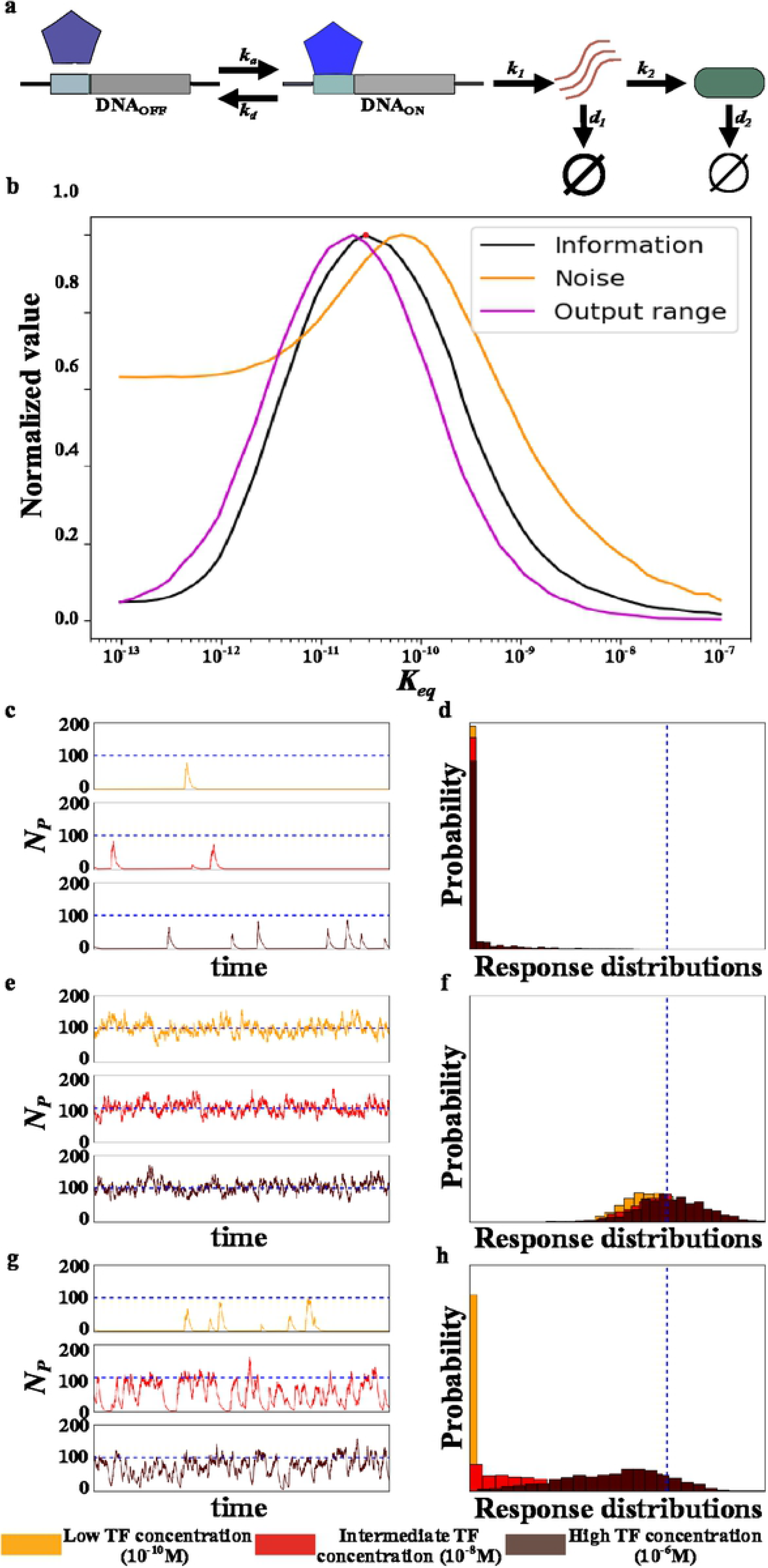
Noise, output range and information in gene regulation. (a) Schematic representation of our model of gene regulation. *k_a_* and *k_d_* correspond to the association and dissociation rate, respectively of a TF with its DNA binding site; *k_1_* and *k_2_* correspond to the mRNA and protein synthesis rate, respectively; *d_1_* and *d_2_* correspond to the mRNA and protein degradation rates, respectively. (b) Information, output range and noise observed in numerical simulations of the system at different TF-DNA affinities (*K_eq_*). Information, noise, and output range are normalized by their respective maximal values. The red circle denotes the affinity at which maximal information is acquired. System behavior at low (*K_eq_*=10^-9^; c and d), intermediate (*K_eq_*=10^-11^; e and f), and high (*K_eq_*=10^-13^; g and h) affinities. Temporal protein dynamics at three different TF concentrations are shown in c, e and g. Response distribution for the same simulations are shown in d, f and h. The blue dashed line in c-h marks the expected mean protein value for constitutive (unregulated, always-on) expression.

We started by analyzing how a TF’s affinity to its *DNA_bs_* affects the relationships between information, output range, and noise (Fig 5b). As in receptor-signal binding, at the lowest affinities, the DNA binding site is almost never bound by TF molecules, regardless of the number of TF molecules (S4a Fig). Hence, little mRNA and protein is produced, independently of the number of TF molecules (Fig 5c and S4 Fig). Response distributions are insensitive to the TF concentration, with a mean close to zero and almost no variation (Fig 5d). For this reason, both noise and output range approach zero as the affinity approaches zero, and so does the acquired information (Fig 5b).

At the highest affinities, noise shows one noticeable difference to the reversible binding of molecules (Fig 2b): it does not decrease to zero (Fig 5b). The reason is that mRNA and protein production have ‘bursty’ dynamics with large excursions from a base line. This bursty dynamics comes from the stochastic nature of mRNA and protein production, which causes fluctuations in the concentration of both kinds of molecules (23). For this reason, gene expression is intrinsically noisy. In particular, for a gene with constitutive expression, the expected number of protein molecules *N_P_* and its expected noise (standard deviation) are equal to *E*(*N_P_*) = (*k_1_/d_2_*)(*k_2_/d_1_*) and 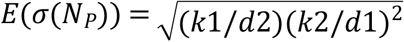 respectively (24). At the highest affinities, the system behaves like a constitutive gene, because TF molecules are almost always bound to the *DNA_bs_* (S4a Fig), and the regulated gene is thus constantly transcribed. Accordingly, in our simulations, the mean number of expressed protein molecules and its standard deviation are close to the expected values for a constitutive gene independently of the number of TF molecules (Fig 5e; S4c Fig; S5 Fig). Because the mean is similar for all number of TF molecules, the output range tends to zero (Fig 5b). However, the amount of noise is higher than zero (Fig 5b) and similar to that expected for a constitutive gene (S5 Fig). As a result, all response distributions are similar to those for the highest concentration of the transcription factor (Fig 5f) and the amount of acquired information about this concentration is small (Fig 5b).

At intermediary affinities, the DNA binding site is not always bound by a TF. It fluctuates between a bound (active) state, when protein molecules are synthesized, and an unbound (inactive) state, when previously synthesized proteins are degraded (Fig 5g, S4a Fig). These fluctuations increase the noise in protein concentrations at intermediate affinities relative to low and high affinities (Fig 5b). They also increase the output range of the system (Fig. 5b). Most importantly, the probability that a binding site is bound by a TF changes with the number of TF molecules, which renders the system’s output – the number of synthesized proteins – sensitive to its input (Fig 5g). Hence, the amount of information acquired about this input increases too (Fig 5b). In other words, noise can increase the acquisition of information also in this gene expression system. Moreover, once again, acquired information is maximal when noise is close to its maximal value (Fig 5b).

### Information, noise and output range in a complete signaling pathway

In a final analysis, we assembled all of the above elements – receptor-signal binding, TF-DNA binding, and gene regulation – into a model of a simple complete signaling pathway. This pathway is akin to a nuclear hormone receptor pathway, such as the signaling pathway of estrogen, progesterone, and various other lipid-soluble signals (31). In this pathway, we quantified the amount of information about the concentration of the input (hormone) signal that is contained in the number of expressed protein molecules. This analysis confirmed our previous results. As in the simpler systems, maximal information acquisition requires high noise, which increases the pathway’s sensitivity to variations in the signal (Supplementary text; S6 Fig).

## Discussion

A fundamental step in signaling pathways is the reversible binding of molecules, which is necessary for the detection of a signal by receptors and for the acquisition of information about this signal. Previous experimental and theoretical work has demonstrated that biological processes, including signaling pathways and their binding interactions, are inherently noisy (23). One would thus expect that the kinetic parameters of binding interactions have evolved to minimize noise, because noise is detrimental for the acquisition of information (13,14,20). However, we find the opposite. The kinetic parameters of signaling pathways must produce noisy binding dynamics or a signaling pathway will acquire little or no information. This is due to the nature of reversible binding interactions. Under biologically sensible parameter values and realistic concentrations of ligands and receptors, binding of molecules is noise-free only when a receptor is completely saturated with its ligand, or if it is unable to bind the ligand. In either case, information acquisition is impossible. Hence, noise in molecular binding is not just unavoidable but necessary for information acquisition in signaling pathways. Importantly, the positive role of noise for information acquisition is not limited to individual binding interactions, but also occurs in more complex systems that include gene expression regulation and more than one binding interaction.

In our models, we observe only one condition where noise is not required for information acquisition. At high signal-receptor affinity, a noise-free ‘perfect’ detection of a signal is possible when the number of receptors is greater than the number of signal molecules. However, producing more receptors than signaling molecules would incur enormous energetic costs. Relatedly, transcriptional regulation generally involves fewer than ten TF binding sites per regulated gene – the analog of a receptor in such a system (29,30) – a number that is much smaller than the average number of transcription factors per cell, which are usually in the hundreds for bacteria and in the thousands for yeast and mammal cells (32,33). Hence, a perfect detection of the number of TFs or signal molecules is not biologically plausible.

Some previous work hinted at a positive role of noise for information acquisition (11,25,27), but this work was not ideally suited to understand the mechanisms by which noise helps increase information acquisition: It did not focus on signaling pathways, did not model molecular interactions explicitly, or it assumed that noise comes from an external source and can be made arbitrarily small. In contrast, our models represent molecular interactions explicitly, which causes noise to emerge naturally from them. In doing so, they also provide a mechanistic explanation of the relationship between noise and information acquisition. However, our models focus on the simplest molecular interactions, and they do not exhaust all possible signaling interactions. Whether other properties of signaling pathways change the way kinetic parameters affect noise and information acquisition is an important task for future work.

Our models include multiple simplifying assumptions. For example, we assumed that the numbers of signaling molecules, receptors, and transcriptional regulators are constant, whereas they may change dynamically in cells. We also considered a simple linear pathway, whereas signaling pathways usually contain regulatory motifs, such as feedback circuits and feed-forward loops (34). In addition, we did not consider molecular interactions such as dimerization (18,19). Similarly, we did not consider the costs of expressing an information processing machinery (20). Because these factors do not affect the nature of reversible binding, we suspect that they might also not reduce the positive role of noisy binding dynamics for information acquisition. However, some of them might increase information acquisition at low noise by other means. For example, some signaling mechanisms increase the amount of information acquired while decreasing noise (11,12,15–19). In contrast to the molecular interactions we study, where noise increases information acquisition by increasing a system’s output range, these mechanisms maintain the output range while decreasing noise (14). It remains to be seen how such different mechanisms interact and jointly affect how biological systems acquire information.

## Methods

### Reversible and consecutive molecular binding models

We consider two kinds of molecules, *S* (signal) and *R* (receptor), which can associate reversibly into receptor-signal complexes at an association rate *k_a_* (M^-1^s^-1^), and a dissociation rate *k_d_* (s^-1^).

To model consecutive reversible binding steps, we assume that, first, a signal (*S*) and a receptor (*R*) reversibly associate into a receptor-signal (*RS*) complex. Second, this complex binds reversibly to a downstream molecule (*D*), such as DNA. We denote the rate of association between the signal and the receptor by *k_aR,S_* (M^-1^s^-1^), and that of dissociation by *k_dRS_* (s^-1^). Similarly, we denote the rate of association between the receptor-signal complex and the downstream molecule by *k_aRS,D_* (M^-1^s^-1^), and that of dissociation by *k_dRSD_* (s^-1^).

### Gene expression system

We model a gene expression system where one chemical species, denoted as *TF* (transcription factor), binds to a DNA binding site (*DNA_bs_*) to regulate the expression of a nearby gene. *TF* molecules associate with the *DNA_bs_* at a rate *k_a_* (M^-1^s^-1^). The dissociation of TF-*DNA_bs_* complexes happens at a rate *k_d_* (s^-1^). In the disassociated state, no transcription occurs, and in the associated state transcription occurs at a rate *k_x_* (s^-1^). Transcribed mRNA molecules are degraded at a rate *d_x_* (s^-1^). Finally, proteins are translated from mRNA molecules at a rate *k_2_* (s^-1^), and degraded at a rate *d_2_* (s^-1^).

### Complete linear signaling pathway

Our model considers the reversible receptor-ligand complex formation and gene expression activation, which is mediated by the receptor-signal complex. Consequently, the parameters that govern the behavior of such a pathway are similar to those described so far, namely: 1) an association rate (*k_aRS_*) between the signal and receptor (R) and a dissociation rate(*k_dRS_*) of the receptor-signal complexes (RS), 2) an association (*k_aRSD_*) and a dissociation (*k_dRSD_*) rate between *RS* and a DNA binding site (*DNA_bs_*), and 3) a rate of gene transcription (mRNA synthesis, *k_1_*), mRNA degradation (*d_1_*), protein synthesis (*k_2_*), and protein degradation (*d_2_*).

### Stochastic simulations

We simulated the behavior of the models described above using Gillespie’s discrete stochastic simulation algorithm (35), using the numpy python package for scientific computing (http://www.numpy.org/). Gillespie’s algorithm captures the stochastic nature of chemical systems. It assumes a well-stirred and thermally equilibrated system with constant volume and temperature. The algorithm requires the probability *p_j_* that a chemical reaction *R_j_* occurs in a given time interval [t,t+τ). Any such probability *p_j_* is proportional to both the reaction rate and the number of reacting molecules. Notice that for first-order reactions, such as the dissociation of a molecular complex into its constituent molecules, *p_j_* is independent of the volume in which the reaction takes place. In contrast, *p_j_* is inversely proportional to the volume in second-order reactions, such as the association of two molecules. For the reversible molecular binding modeled here, the probabilities *p_a_* and *p_d_* that two molecules associate and dissociate, respectively, are proportional to

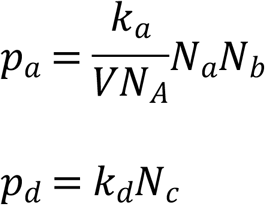

where *V* is the reaction volume, *N_A_* is Avogadro’s number, and *N_a_, N_b_*, and *N_c_* are the numbers of molecules of the two chemical species *a* and *b* and of the complex *c*.

The probabilities *p_mRs_, p_mRd_, p_Ps_* and *p_Pd_* of a mRNA transcription, mRNA degradation, protein synthesis, and protein degradation event are given by

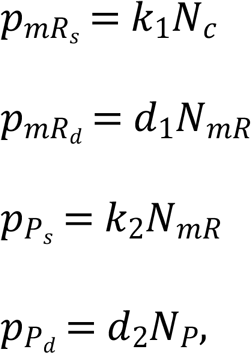

respectively. In these expressions, *t*he quantities *N_mR_* and *N_P_* are the numbers of mRNA molecules and of protein molecules, respectively. We model a haploid organism with only a single DNA binding site, corresponding to a single regulated gene. In this case, the probability of mRNA synthesis can be reduced to

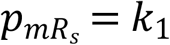

when the binding site is bound by transcription factor (*N_c_*=1) and to

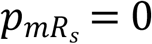

when the binding site is unbound (*N_c_*=0).

### Initial conditions for simulations

To determine the initial conditions of the system, we first calculated the expected number of complexes formed as

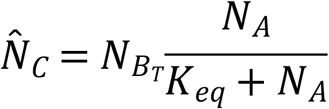

The quantity *N_A_* is the number of signal or TF molecules and *N_Bt_* is the total number of receptor molecules or DNA binding sites. Notice that 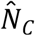 is a real number, and for the specific case of the TF-*DNA_bs_* interaction it can only take a value between 0 and 1. Thus, 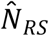 equals to the probability that the DNA binding site is bound by a transcription factor. However, for the receptor signal complexes, it represents the number of complexes formed. We selected the initial state of the number of complexes (*N_C_i__*) at random with binomial probability for the binding site. However, we selected the closest integer to 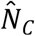 for the receptor signal case. Then we defined

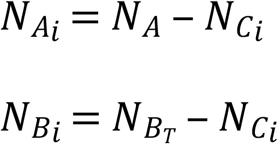

as the initial state of the number of free signal or TF molecules (*N_A_i__*) and of the receptors or binding sites (*N_B_i__*). Finally, as the initial state of the number of mRNA and protein molecules we used

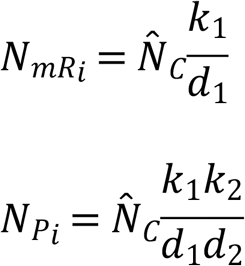

which is the expected average number of mRNA and protein molecules for a constitutively expressed gene (23), multiplied by the probability that the DNA binding site is bound by a TF molecule.

### Information quantification

The number of molecules of any chemical species in a cell or in a unit volume fluctuates, because molecules are produced and decay stochastically, and because they undergo random Brownian motion caused by thermal vibrations. We use Shannon’s entropy to quantify the unpredictability caused by such stochastic fluctuations in a signal as

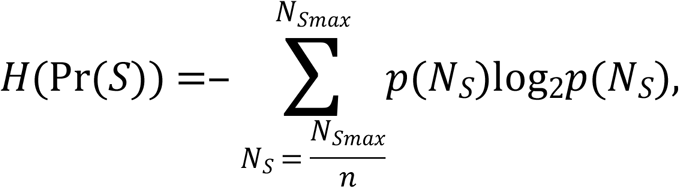

where Pr(S) is the probability distribution of the signal, and *p*(*N_S_*) is the probability that the system contains *N* molecules of the signal. In our models the signal represents either a molecular signal or cue (2).

For all our analyses, we performed at least 1000 simulations for each of *n* different numbers of signal molecules that were evenly distributed within the interval [*N_Smax_/n,N_Smax_*] (*n*≤*N_Smax_*). For this reason

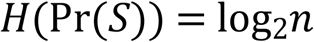

Signals trigger changes in a cell’s state that produce a response or output (*O*) of the system, such as the production of molecules. A cell acquires information when the output *O* reflects (fully or partially) the value of S. This information can be quantified via the mutual information:

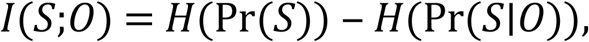

which is equal to the entropy *H*(Pr(*S*)) minus the conditional entropy *H*(Pr(*S|O*)), which represents the entropy of *S* given that *O* is known (13). In other words, the mutual information *I* quantifies the acquired information as the amount of information that an output chemical species *O* harbors about a signal *S*.

### Noise quantification and output range

The systems we model produce a probabilistic response for any given quantity *N_S_* of the signal. This response can thus be represented as a conditional probability distribution:

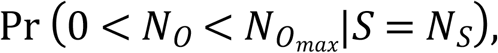

where *N_O_* and *N_Omax_* are the number and maximal number of output molecules, respectively. We estimated this response distribution through at least 1000 replicate simulations of the system for each value of *N_S_*. We then quantified noise as the standard deviation of the response distributions, averaged over all *n* possible values of *N_S_*:

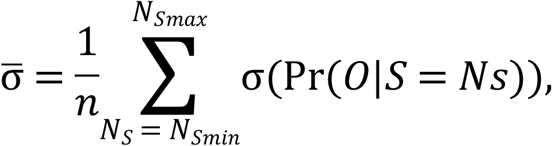

We define the output range as the difference between the maximal and minimal mean value of all response distributions.

### Parameter values

Our simulations considered the following biologically sensible parameter ranges. The association and dissociation constants *k_a_* and *k_d_* of reversible complex formation define the equilibrium constant *K_eq_*=*k_d_/k_a_* (M), which we used in our simulations. The smaller *K_eq_* becomes, the more association becomes favored over dissociation (28). In particular, for the binding between ligands and (nuclear) receptors, we used values of *K_eq_* within the interval [10^-6^M,10^-6^M], because the micromolar to nanomolar range is common for such complexes (28,32,36–38). For TF-DNA binding, empirical data suggests that usually *K_eq_*<10^-8^ and can reach picomolar (10^-12^M) or even smaller values (28,32,36–38). Thus, we used values in the interval [10^-8^M,10^-12^M].

For mRNA, experimentally measured half-lives usually lie in the range of seconds to hours (39–42). Protein half-lives typically lie between hours and days (41,43). Taking all this information into consideration, we chose mRNA half-lives within the interval [1min,30min], and protein half-lives where within [15min, 3h]. We assume that the ratio *k_2_/k_1_*, which describes the speed of the protein synthesis rate relative to the mRNA synthesis rate, exceeds 1.0 (44). Because the residence time of transcription factors on DNA lies within seconds to hours (45,46), we assumed a residence time within this interval [10sec,2h]

Finally, we always considered concentrations of molecules to lie within the interval [10^-9^M,10^-6^M], because these are typical concentration of most molecules within a cell or a nucleus (47). Notice that for some of our simulation we also needed to explore values outside these ranges. Specific parameter values used for each simulation are listed in supplementary tables 1-4.

## Acknowledgments

We acknowledge support from the European Research Council under Grant Agreement No. 739874, by Swiss National Science Foundation grant 31003A_172887, as well as by the University Priority Research Program in Evolutionary Biology.

## Supporting information captions

**S1 Fig.** Mean number of receptor-signal complexes 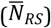 formed at different affinity values (*K_eq_*). The maximum number of receptor-signal complexes for this simulation is 50 (see S1 Table).

**S2 Fig.** Noise, output range and information observed in numerical simulations of the receptor-signal system at different affinities (K_eq_) and with different concentrations of *R* and *S*. a) *R*=10^-9^M and *S*=[10^-9^M,10^-7^M]. b) *R*=10^-8^M and *S*=[10^-8^M,10^-6^M]. a) *R*=10^-7^M and *S*=[10^-7^M,10^-5^M]. Information, noise, and output range are normalized by their respective maximal values.

**S3 Fig. Information and noise in two consecutive binding interactions with ten DNA_bs_.** Contour plots of (a and d) noise, output range (b and e) and information acquisition (c and f) in the receptor-signal complex (*RS*; a-c) and in the receptor-signal-*DNA_bs_* complex (*RSD*; d-f) as a function of the affinities between both the receptor and the signal (*K_eqR,S_*), and the receptor-signal complex with the downstream molecule (*K_eqRS,D_*). Red-dashed rectangles circumscribe biologically sensible receptor-signal DNA affinities ([10^-8^M,10^-13^M]) and receptor signal affinities ([10^-6^M,10^-9^M]). White-dashed rectangles delineate the region of maximal information acquisition at the receptor-signal-*DNA_bs_* level. Acquired information, output range and noise are plotted from minimally to maximally observed values, color-coded as indicated by the color bar.

**S4 Fig.** (a) Probability of TF-DNA binding, (b) mean number of mRNA molecules 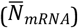, and (c) mean number of protein molecules 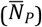, as a function of TF-DNA affinity (*K_eq_*). For the parameters we used (S3 Table), the expected mean number of mRNA and protein molecules produced from a constitutively expressed gene is 2 and 100, respectively.

**S5 Fig. Non-normalized values of noise in the number of proteins.** The black line is the standard deviation expected for the parameters used in this simulation (S3 Table).

**S6 Fig. Contour plot of acquired information and noise in a complete signaling pathway.** (a) Information and (b) noise for different affinities between receptor and signal molecules (*K_eqR,S_, y* axis), and receptor-signal complexes with a DNA binding site (*K_eqRS,D_, x* axis) in a model of a simple lineal signaling pathway where a signal bound receptor can bind DNA and regulate gene expression. The number of signal molecules constitute the input into this pathway, and the number of protein molecules expressed from the regulated gene constitute the output. The red-dashed rectangle show experimentally measured affinity values between receptors and signals ([10^-6^M,10^-9^M]) and between transcriptional regulators and DNA ([10^-8^M,10^-13^M]). The amount of acquired information and noise level are indicated in the color bar.

